# Uukuniemi virus infection causes a pervasive remodelling of the RNA-binding proteome in tick cells

**DOI:** 10.1101/2024.09.06.611624

**Authors:** Alexandra Wilson, Wael Kamel, Kelsey Davies, Zaydah R. De Laurent, Rozeena Arif, Lesley Bell-Sakyi, Douglas Lamont, Yana Demyanenko, Marko Noerenberg, Alain Kohl, Shabaz Mohammed, Alfredo Castello, Benjamin Brennan

## Abstract

Cellular RNA-binding proteins (RBPs) are pivotal for the viral lifecycle, mediating key host-virus interactions that promote or repress virus infection. While these interactions have been largely studied in the vertebrate host, no comprehensive analyses of protein-RNA interactions occurring in cells of arbovirus vectors, in particular ticks, have been performed to date. Here we systematically identified the responses of the RNA-binding proteome (RBPome) to infection with a prototype bunyavirus (Uukuniemi virus; UUKV) in tick cells and discovered changes in RNA-binding activity for 283 proteins. In an orthogonal approach, we analysed the composition of the viral ribonucleoprotein by immunoprecipitation of UUKV nucleocapsid protein (N) in infected cells. We found many tick RBPs that are regulated by UUKV infection and associate with viral nucleocapsid protein complexes. We confirmed experimentally that these RBPs impact UUKV infection. This includes the tick homolog of topoisomerase 3B (TOP3B), a protein able to manipulate the topology of RNA, which showed an effect on viral particle production. Our data thus reveals the first protein-RNA interaction map for infected tick cells.

**Research highlights:**

- UUKV RNAs interact with nearly three hundred tick cell RBPs.
- Demonstrated an enrichment of N protein interactors within the upregulated RIC data suggesting a direct involvement in viral RNA metabolism and translation.
- Developed a robust methodology to silence gene expression in tick cell cultures.
- The TOP3B complex facilitates efficient packaging of UUKV virions.

## INTRODUCTION

*Uukuvirus uukuniemiense* is the prototypic virus within the genus *Uukuvirus* of the family *Phenuiviridae* that was first isolated from *Ixodes ricinus* ticks in Finland in 1964 [1]. Uukuniemi virus (UUKV; strain S-23) has been utilised as the prototype tickborne bunyavirus for several decades and has contributed to many aspects of bunyavirus research [2,3], such as the determination of the tri-segmented nature of bunyaviruses [4], the isolation of the first RNA-dependent RNA polymerase from a bunyavirus [5], the determination of the structural composition of the virion [6] and entry into mammalian cells [7–9]. Like other phenuiviruses, the genome of UUKV comprises three segments of negative or ambi-sense RNA, named small (S), medium (M) and large (L) which are deposited into the cytoplasm of infected cells as ribonucleoprotein (RNP) complexes encapsidated by the viral nucleocapsid (N) protein and associated with the viral RNA-dependent RNA polymerase (L protein). Both the N and L proteins can interact with and bind viral RNA, although currently it is unclear if this binding capability extends to cellular RNA [10,11]. The N protein has also been shown to interact with the L protein, the viral glycoproteins and a range of cellular proteins during infection [12,13]. The virus also encodes a non-structural protein (NSs) within the S segment that has been demonstrated to be a weak interferon antagonist [14]. However, unlike other phenuiviruses transmitted by mosquitoes or sandflies, the M segment of tickborne phenuiviruses only encodes the glycoprotein precursor and does not encode any other non-structural proteins [15].

Research into tickborne bunyaviruses has been heavily biased towards studies in vertebrate systems, primarily due to the expertise needed and expense of facilities associated with working with live ticks and the lack of genomic data or molecular reagents to conduct virus infection experiments within tick-derived cell lines. To date, over 70 cell lines have been developed from multiple tick species of medical and veterinary importance [16]. However, only within recent years have detailed whole tick genomes and the genome of an *Ixodes scapularis* cell line been sufficiently annotated to support studies into the molecular interactions of tickborne pathogens with host cells [17–19]. These tools have facilitated several groups to conduct proteomic, transcriptomic and genomic analyses of tick-derived cell lines [20–28]. However, no research has been carried out to determine the host proteins with pivotal roles in the lifecycle of a bunyavirus within tick cells.

Viruses, including those with a RNA based genome, have a limited coding capacity, and therefore cannot encode all the proteins required for a fully autonomous lifecycle, relying on host resources to replicate and spread. For example, viruses are fully dependent on the translation apparatus of the host cell to synthesise viral proteins. However, protein synthesis is one of the steps of virus infection, and cellular RNA binding proteins (RBPs) are expected to additionally participate in RNA stability, replication and packaging of the viral RNA within the viral particles [29]. RBPs are also critical in the cellular defence against viruses, and in vertebrates, virus sensors and effectors recognise molecular patterns that are specific to viral RNAs. In invertebrates, it is believed that the RNA interference (RNAi) pathway, which utilises many RBPs such as Dicer and Argonaute proteins, is the main effector of the antiviral response. However, whether other antiviral mechanisms involving RBPs exist in arthropods, as demonstrated in vertebrates, remains unexplored. Despite the central roles that RNA plays in the viral lifecycle, the complement of host RBPs that participate in arboviral infection remain largely unknown, particularly for arthropod vectors such as ticks [30–33]. This has led to a dearth of information and insight as to why and how these important vectors can cause consequential diseases globally.

In this manuscript, we utilise our proteome-wide approach called RNA interactome capture (RIC) to study the responses of the tick RNA-binding proteome (RBPome) of the *Ixodes scapularis* cell line ISE6 to infection with UUKV [34,35]. We discovered that the tick RBPome is plastic and reconfigures in response to infection, with nearly three hundred proteins exhibiting either an increased or decreased RNA-binding activity. In addition, we conducted an orthogonal approach to isolate the viral ribonucleoproteins via immunoprecipitation of the UUKV N protein. Strikingly, we found 18 RBPs with virus-regulated RNA-binding activity that are associated with the viral ribonucleoproteins. We validated our findings with dsRNA knockdowns in ISE6 cell cultures showing that the discovered RBPs do regulate virus infection. Initial characterisation of the selected candidates revealed a role for the protein ISCI010954 (TOP3B in vertebrates) in the production of infectious virus particles by facilitating the efficient packaging of UUKV RNAs into virions.

## RESULTS

### Establishing comparative RNA interactome capture (cRIC) in tick cells

Earlier studies showed that virus infection causes a pervasive remodelling of the cellular RBPome in human cells [31,32,36]. Our goal was to determine whether such a phenomenon also occurs during infection of cells from the tick, and if so, whether it is biologically relevant. We then used the established conditions to perform an experiment comparing uninfected and UUKV-infected cells [34] (Fig 1A). To determine which timepoint to utilise for this analysis, we conducted a viral growth curve to examine UUKV infection kinetics in ISE6 cells. Release of infectious virus was detected throughout the course of infection and progressively increased until the final time point at 12 days post infection (p.i.) (Fig 1B). In parallel, expression of the viral nucleocapsid protein N was monitored in infected cell monolayers. UUKV N protein was only detectable by Western blotting from 9 days p.i. (Fig 1C). We selected 9 days p.i., for the analysis because there was a strong increase in UUKV N abundance from 9 to 12 days p.i., which implied active replication. UUKV-infected cell monolayers were also analysed by confocal microscopy through staining the monolayer for the presence of the UUKV N protein. This qualitative assessment (Fig 1D) provided assurance that most cells were infected at this timepoint and confirmed its suitability for cRIC analysis (Fig 1D).

**Fig 1.**
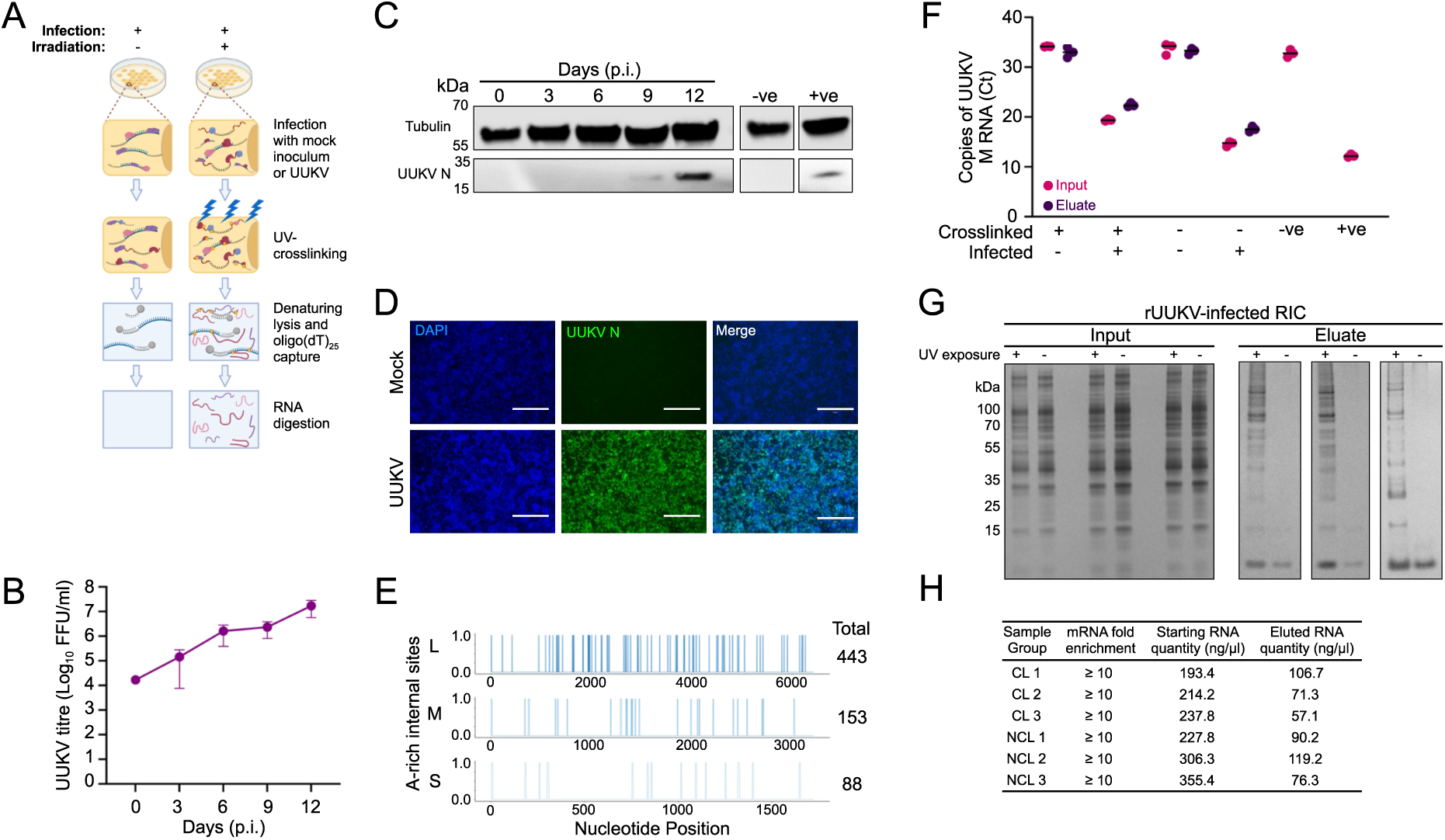
Preparation of mock and UUKV-infected RNA interactome capture (RIC) samples derived from ISE6 tick cell cultures. (A) Schematic showing the methodology of interactome capture utilising UV-crosslinking and oligo(dT)_25_ capture beads. (B) Titre of UUKV in supernatant of infected ISE6 cells up to 12 days p.i. Data are plotted as the mean virus titre (FFU/ml) ± SD of n=3 biological replicates. (C) Representative images of cell extracts from cell monolayers in (B) probed with anti-α Tubulin, and anti-UUKV N antibodies. ‘-ve’: mock-infected cell lysate; ‘+ve’: sample known to be infected with UUKV derived from infected mammalian cells. (D) UUKV-infected ISE6 cell monolayers stained with DAPI (blue), and probed for UUKV N protein (green) at 9 days p.i. Cell monolayers were imaged using an EVOS microscope using a 10x objective. Scale bar is equal to 300 μm. (E) The poly(A) adjacent content of the UUKV genome segments. The involvement of each nucleotide in poly(A)/poly(A)-like sites is visualized. (F) Quantity of UUKV M RNA within input and eluate samples of ISE6 cell monolayers treated by RIC. (G) Proteins found in UUKV-infected ISE6 cells, derived from input or eluate RIC samples, visualised via silver staining. (H) Properties of RNA found within the samples described in (G). RT qPCR was performed against the 18S ribosomal RNA and eukaryotic elongation factor 1-alpha (*ELF1A*; mRNA used as a housekeeping gene) The ΔΔCT method was used to measure the quantity of 18S RNA normalized to *ELF1A* RNA levels. The mRNA fold enrichment in the elute compared to the input sample was calculated for each biological sample. Finally, the RNA quantities in the samples from (G) were measured via a Nanodrop spectrophotometer.

During initial infection, the ratio of viral RNA to cellular mRNA increases and this can lead to interference in cellular mRNA production [37]. To obtain a complete picture of the intracellular environment during UUKV infection of ISE6 cells, it was therefore important to ensure the capture of both cellular and viral RNA. This presents a problem, as bunyaviral genomic RNA or mRNAs are non-polyadenylated [38,39] and the oligo(dT)_25_ capture beads used in cRIC exploit the interaction between the poly-A tail of mRNAs. We analysed the genome of UUKV for the presence of ‘poly-A like’ regions, which we defined as a minimum of five sequential A nucleotides and any larger region containing a ≥80% A content. Our hypothesis was that oligo(dT)_25_ capture beads would be able to capture the UUKV RNA by interacting with these ‘poly-A like’ regions, allowing the interactome capture to reflect both viral and cellular RNA within the infected cells. Our *in-silico* analysis revealed 88, 153 and 443 poly-A-like regions present within the UUKV S, M or L RNAs, respectively (Fig 1E). These results provided a basis for continuing with the UUKV-infected cRIC experimental protocol. Recent data utilising Rift Valley fever virus (RVFV) has shown that while the majority of viral RNAs are encapsidated by the N protein, some regions of the viral genome/antigenome RNAs and the subgenomic mRNAs remain exposed, which may facilitate RBP-binding [40].

We next assessed the isolation of UUKV RNAs using RT-qPCR in cRIC eluates focusing on the M segment. UUKV M RNA was detected in all infected samples but not in the mock samples (Fig 1F). These data confirmed that oligo(dT)_25_ can isolate UUKV transcripts (Fig 1E & 1F). In parallel, cRIC inputs and eluates were tested for suitability for subsequent protein analysis. Silver staining revealed a discrete protein pattern in eluates of UV-irradiated samples that was consistent with similar experiments done in human and fruit fly cells (Fig 1G) [41]. This pattern was different from that of the whole cell lysate (inputs of the cRIC experiment), indicating that these isolated proteins were a subset of the cellular proteome. No proteins were detected in non-irradiated samples, further confirming that the isolated proteins were RBPs.

### UUKV infection induces a reconfiguration of the tick RBPome

We next performed a quantitative proteomics analysis of the cRIC eluates. Quality control analyses revealed that i) the overall protein intensity in UV cross-linked samples is far superior to that in non-irradiated counterparts (Fig S1A); and ii) that the replicates clustered together in Principal Component Analysis (PCA) with UV irradiation explaining most of the variation (∼85%) followed by whether cells were infected with UUKV (∼6%) (Fig S1B). From these results it was established that data were of excellent quality and could be used for further analyses.

From the 572 identified proteins, 541 and 530 were enriched in UV-irradiated over non-irradiated samples at a false discovery rate (FDR) of 10% and 1%, respectively (Fig 2A). Due to the limited available information and annotation for ticks, we decided to include the full 10% FDR group for further analyses. To test if these proteins were RBPs, we first identified the human *(Homo sapiens*) and fruit fly *(Drosophila melanogaster*) orthologous proteins using InParanoid [37]; we found that 65% and 58% of the proteins within our dataset had human or fruit fly orthologs, respectively (Fig. 2B). Approximately 80% of the 349 proteins with human orthologs have been experimentally determined to be RBPs (Fig. 2C). Therefore, we concluded that the ISE6 tick RNA interactome is consistent with a *bona fide* RBPome.

**Fig 2.**
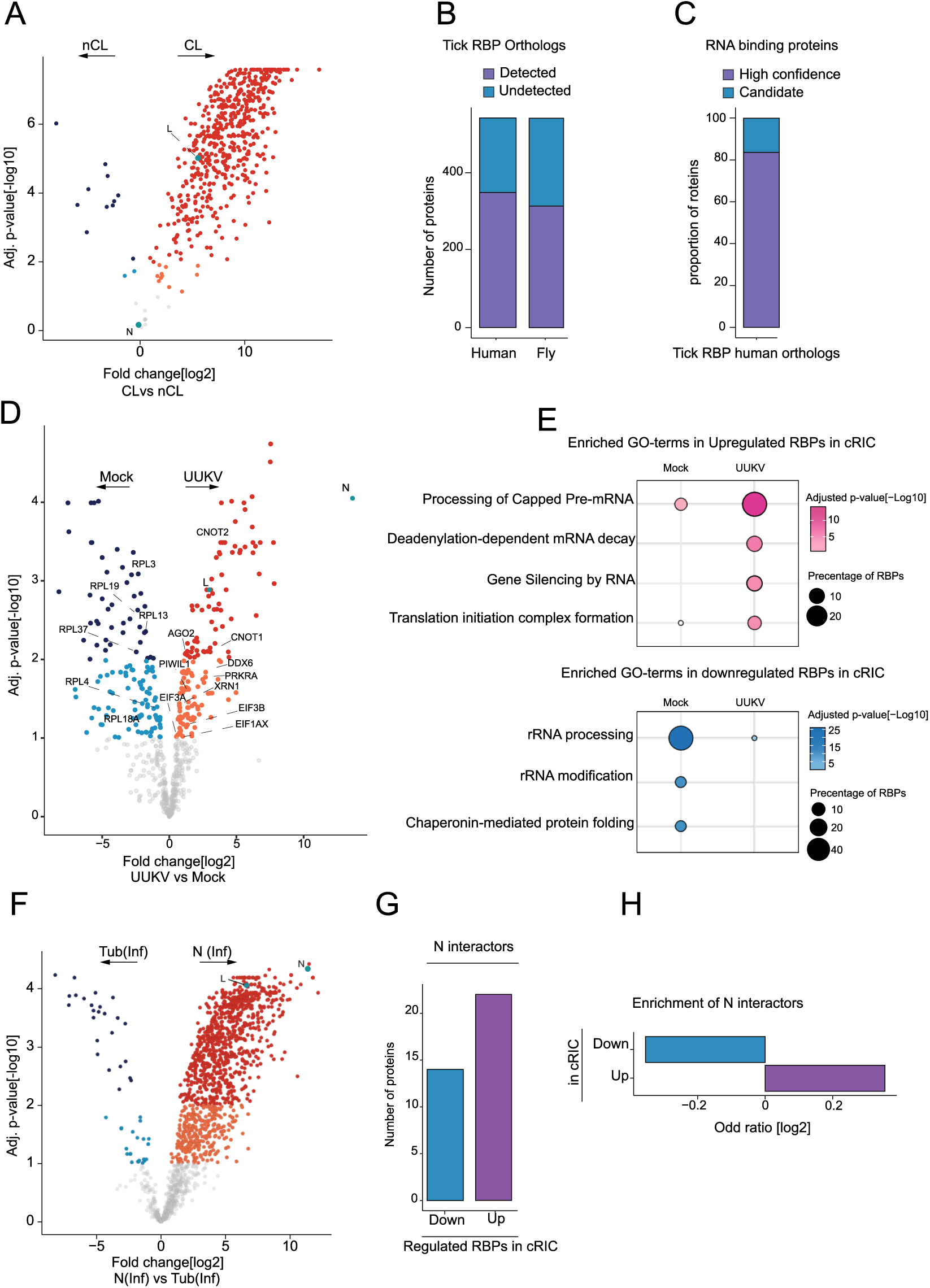
Differential regulation of the RBPome during UUKV-infection in ISE6 cells. (A) The fold changes in the proteins found in the UUKV-infected cross-linked samples compared to the non-cross-linked samples (x-axis) were plotted against the significance of this fold change (Adj. p value[-log10], y axis). For proteins with a positive fold change, proteins at a 1-10% false discovery rate are coloured in orange and proteins at a 1% false discovery rate or below are coloured red. For proteins with a negative fold change, the false discovery rate groups are coloured light blue (1-10% FDR) and dark blue (1% FDR), respectively. (B) The numbers of proteins within the differential ISE6 RBPome with corresponding human and *Drosophila* (fly) orthologs. (C) The human orthologs identified in (B) were compared against a defined experimentally proven database of RNA binding proteins (RBPbase v0.2.1 alpha). Proteins which were found by three independent comparisons were termed high confidence (purple), while the other proteins were termed candidates. (D) The fold changes in the proteins identified in (A) were compared to the mock cross-linked samples (x axis) and plotted against the significance of this fold change (Adj. p value[-log10], y axis). For proteins with a positive fold change, proteins at a 1-10% false discovery rate) are coloured in orange and proteins at a 1% false discovery rate or below are coloured red. For proteins with a negative fold change, and therefore in higher quantities in non-cross-linked samples, proteins at a 1-10% FDR coloured in light blue, and proteins at a 1% FDR or below are coloured dark blue. Proteins with greater than 10% FDR are shown in grey. (E) Enrichment of cellular pathways in the upregulated (shown in pink) and downregulated (shown in blue) RNA binding proteins in the cRIC during UUKV-infection. (F) Comparison of fold changes of proteins found in the UUKV-infected samples using UUKV N or tubulin antibody immunoprecipitation. (G) The number of differentially regulated RBPs in cRIC interacting with the viral nucleocapsid (N) protein. (H) The odds ratio of the N interactors found in the upregulated and downregulated RIC RBPs were calculated to determine the enrichment of the N interactors in the RIC data.

Strikingly, 283 RBPs were differentially regulated in infected versus mock-infected cells at a 10% FDR (Fig. 2D). Reassuringly, both the RNA-dependent RNA polymerase (L) and the viral nucleocapsid protein (N) were amongst the most enriched proteins in UUKV-infected cells compared to mock-infected samples, with N levels exhibiting the highest fold change observed. Gene set enrichment analysis revealed that RNAi silencing, capping, deadenylation and translation were the most upregulated pathways, while rRNA processing was depleted under infection conditions. Our results are consistent with the known importance of RNAi in the antiviral response of arthropods and translation in viral protein synthesis [42,43].

### The UUKV N interactome and its connections with RBPome dynamics

Bunyaviral RNAs are found strongly associated with the viral nucleoprotein (N), which is one of the main components of the viral ribonucleoprotein (vRNP) complexes present in the cellular cytoplasm [44]. To survey the range of proteins associated with bunyaviral RNAs, we investigated the N protein interactome in UUKV-infected tick cells by using immunoprecipitation (IP) with highly specific antibodies. The proteomic data correlated well, and experimental triplicates clustered together in PCA analysis with 87.4% of the variance explained by infection status and 7.1% by the antibody used (i.e. against N or α-Tubulin) (Fig S1). A total of 701 and 922 proteins were enriched in N over β-Tubulin IP conditions with 1% and 10% FDR, respectively (Fig 2F). In addition, 613 proteins were enriched in UUKV-infected cells over mock conditions at 10% FDR (Fig. S1D). Both N protein and the RNA-dependent RNA polymerase (L) were enriched in infected conditions and over the β-Tubulin control IP. For a more stringent analysis, we cross-referenced the 10% FDR candidates described in the two different comparisons above (Fig. 2F and S1D). A total of 378 proteins were consistently enriched in UUKV-infected vs mock and N vs β-Tubulin IP conditions (Fig. S1E), and these were classified as N interactors. We cross-referenced the N interactors with our previously generated differential RBPome (defined in Fig 2D). Of the 378 proteins immunoprecipitated from UUKV-infected cells, 14 and 22 proteins were found within the downregulated and upregulated RBP groups, respectively (Fig. 2G). Interestingly, we found a modest enrichment of N interactors within the upregulated RIC data suggesting their direct involvement in vRNA metabolism (Fig. 2H).

### dsRNA knockdown of cellular RBPs regulates UUKV infection

To evaluate the roles of the UUKV-responsive RBPs and their importance in tick cell infection, we knocked down several candidates using dsRNA transfection (Fig 3). We utilised a previously optimised Magnetofection technology to deliver dsRNA into tick cells [45]. After dsRNA transfection, various parameters such as cell metabolic activity, UUKV N expression, intra- and extracellular viral RNA (RT-qPCR); and release of infectious virions (IFA analysis) were measured. Confocal microscopy analysis of UUKV-infected ISE6 monolayers showed a robust UUKV infection over the time course in both the mock- and dseGFP transfected monolayers. In contrast, transfection of a dsRNA targeting UUKV N almost completely abrogated UUKV infection (Fig 3A). No difference in UUKV RNA abundance, N protein synthesis or viral particle production was observed in dseGFP-transfected monolayers compared to the mock-transfected samples (Fig 3B). Conversely, a significant decrease in all measurements was observed after transfection with dsRNA targeting the UUKV N coding region (Fig 3C). Following transfection with either dsRNA, only small variations in cell metabolic activity were observed. These data validated our dsRNA delivery approach into tick cell cultures.

**Fig 3.**
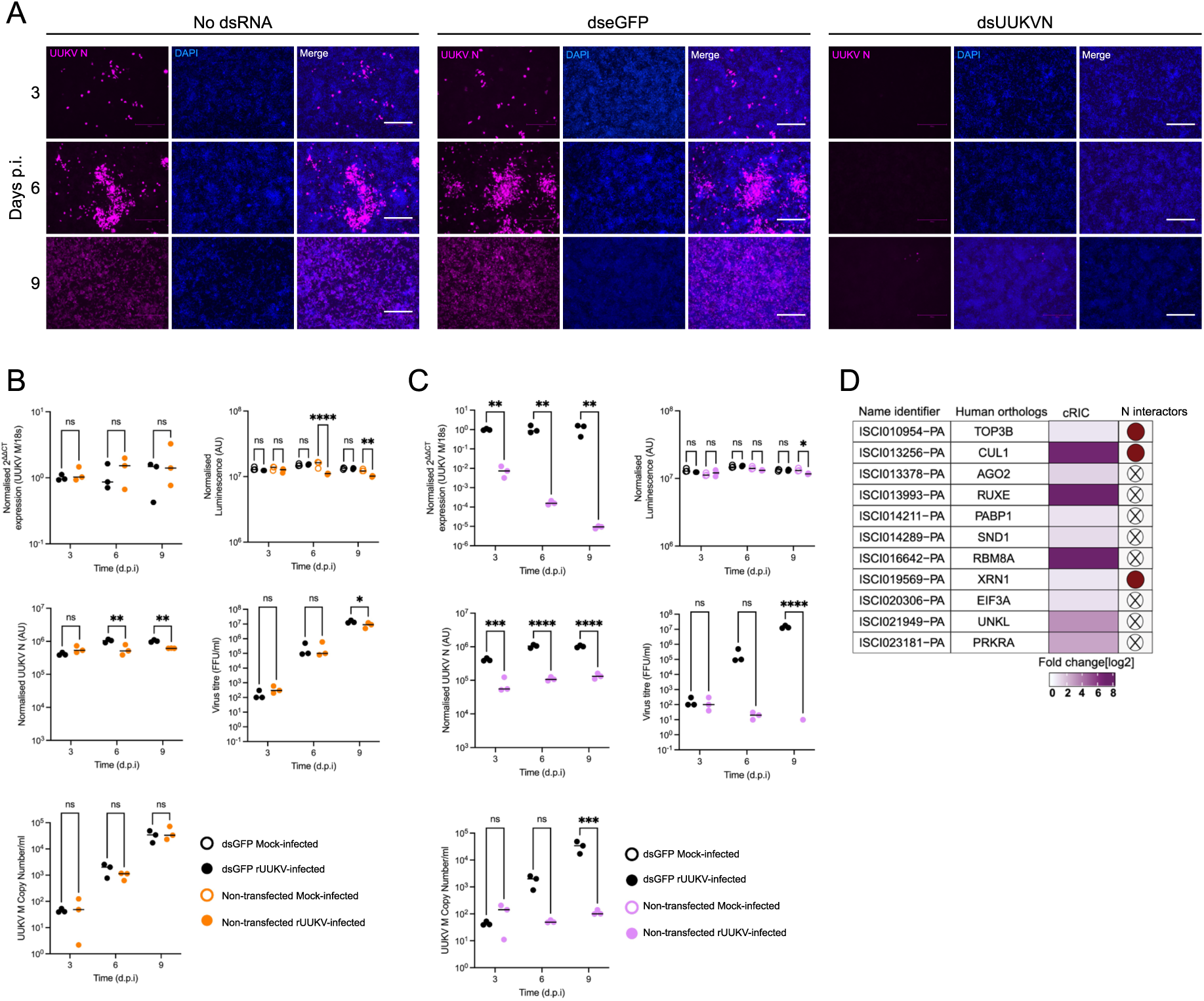
Effect of eGFP dsRNA or UUKV N dsRNA transfection on UUKV replication in infected ISE6 cells. (A) UUKV-infected, dsRNA-transfected cell monolayers, imaged at the time points indicated were stained using DAPI (blue) and mouse anti-UUKV N (purple). Imaging was carried out using an EVOS microscope, scale bar 300 μm. In parallel, cell monolayers were either transfected as (B) mock (black) vs dseGFP (orange) or (C) mock (black) vs dsUUKV N (pink) and were further analysed for additional parameters. These include quantity of intracellular UUKV M RNA (normalised 2^ΔΔCT^), cell metabolic activity (normalised luminescence), UUKV N expression via in well western blot (normalised UUKV N), UUKV titre (FFU/ml) and UUKV M RNA (UUKV M RNA copy number/ml) present in cell culture supernatant. Each gene type knockdown biological replicate was carried out conjointly with all other gene type knockdowns alongside the positive and negative controls. (D) Targets were selected for knockdown analysis and their corresponding human orthologs. The fold change within the cRIC data [log2] is indicated in purple as defined in Fig. 2D, alongside whether the protein is defined as an NCAP interactor as specified in Fig. 2G. Statistical significance was measured by ordinary two-way ANOVA with Tukey’s multiple comparisons test. Asterisks indicates significance **** = p < 0.0001, *** = p ≤ 0.001, ** = p ≤ 0.01, * = p ≤ 0.05, ns = not significant.

To assess the impact of the UUKV-responsive RBPs, we used the optimised protocol with dsRNAs against the mRNAs encoding our candidate RBPs or eGFP as control. Most of the targeted transcripts were effectively depleted in transfected cells apart from those of the *CUL1*, *PRKRA*, and *XRN1* genes that showed only 50% knockdown efficiency at late timepoints (Fig S2). Commercially available antibodies against tick proteins are currently lacking and, therefore, RNA analysis by RT-qPCR is the only reliable method to assess gene silencing. We also examined cell metabolic activity and found no effect over the eGFP control for most of the dsRNAs over the time course. However, a reduction in metabolic activity was observed in UUKV-infected samples at several timepoints for knockdowns targeting PABP1, PRKRA and UNKL proteins (Fig S3). The potential effects of altered viral fitness will be discussed below.

Many of the RBP knockdowns resulted in a statistically significant increase in UUKV M segment RNA at either the 3- or 6-day time point (Fig 4). This is not unexpected considering the known antiviral roles of proteins such as AGO2 and XRN1 in other mammalian and arthropod systems [21,46–48]. The knockdown of other genes involved in the translation of cellular mRNAs such as EIF3A and PABP1 also resulted in an increased accumulation of UUKV mRNA in the cell. Conversely, TOP3B knockdown had no impact on UUKV M RNA levels (Fig 4).

**Fig 4.**
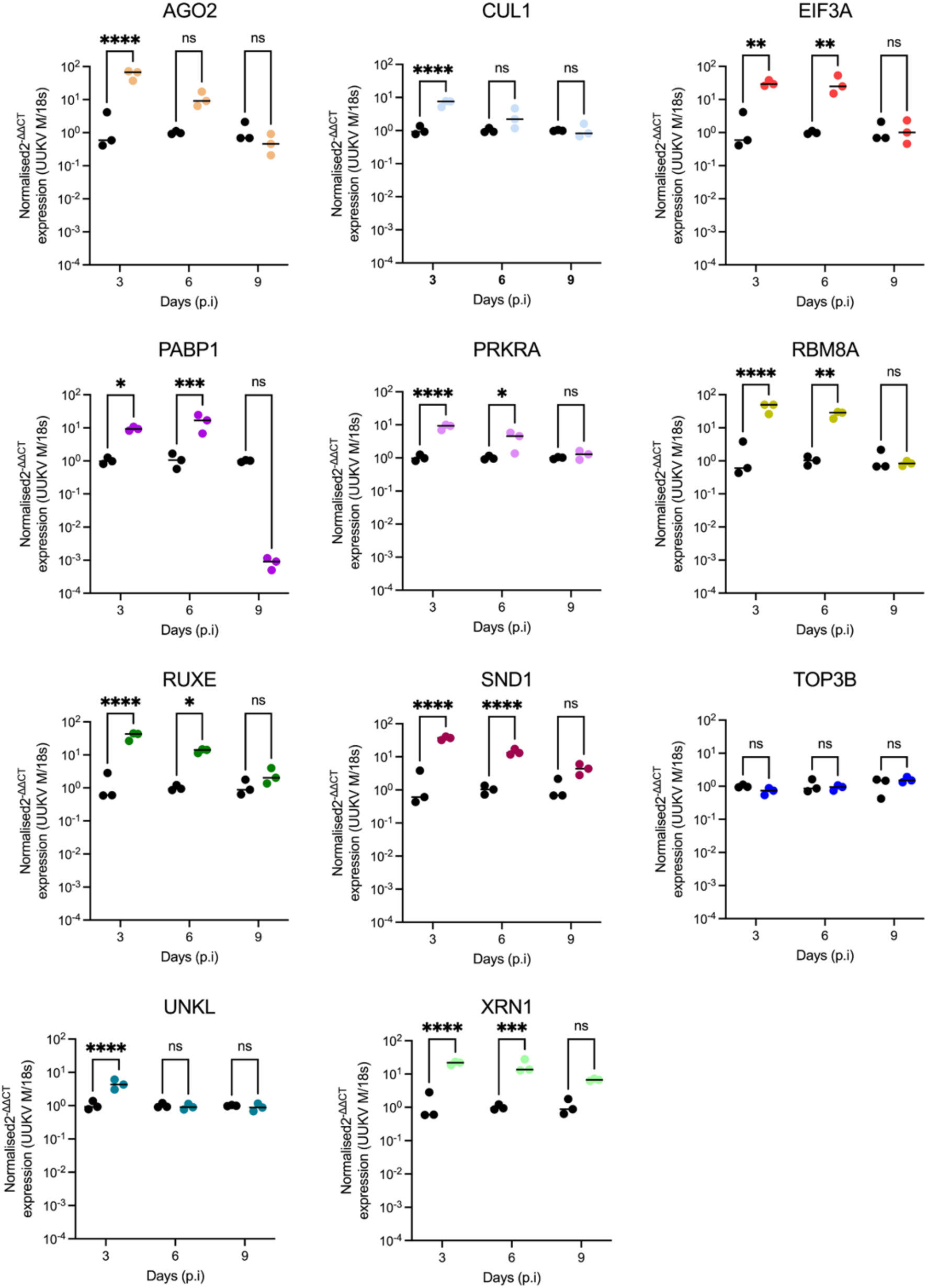
Effect of dsRNA transfections on the expression of UUKV M segment RNA levels within UUKV infected ISE6 cell culture. ISE6 cell monolayers were transfected with 2µg of dsRNA homologous to the indicated gene, before being infected with UUKV at an MOI of 5 in triplicate and harvested at the indicated timepoints. Data presented show the normalised expression of UUKV M RNA in UUKV-infected, dsRNA-transfected ISE6 cell monolayers, calculated using the 2-ΔΔCt method. Black symbols represent dseGFP data (mock) and coloured symbols represent dsGene data. Statistical significance was measured by ordinary two-way ANOVA with Tukey’s multiple comparisons test. Asterisks indicates significance **** = p < 0.0001, *** = p ≤ 0.001, ** = p ≤ 0.01, * = p ≤ 0.05, ns = not significant.

Both the release of infectious virus particles (Fig 5) and the presence of viral RNA in the supernatant (Fig 6) were assessed over the infection time course. Only a small amount of infectious virus was detectable in the supernatant at 3 days p.i. (∼ 10^2^ FFU/ml) in the dseGFP treated control. This increased over time to ∼10^6^ FFU/ml by 9 days p.i. Surprisingly, the increased detection of UUKV M RNA in the knockdown cells observed in Fig 4 did not translate into an increase of infectious viral particles at the time points tested. An observable and significant decrease in the production of infectious UUKV was observed at 9 days p.i. in all knockdown samples except for UNKL and XRN1, in which no significant difference was observed (Fig 5). This decrease is likely due to a reduction of cell metabolic activity, perhaps because of the observed increase in viral fitness (M RNA levels) (Fig S2).

**Fig 5.**
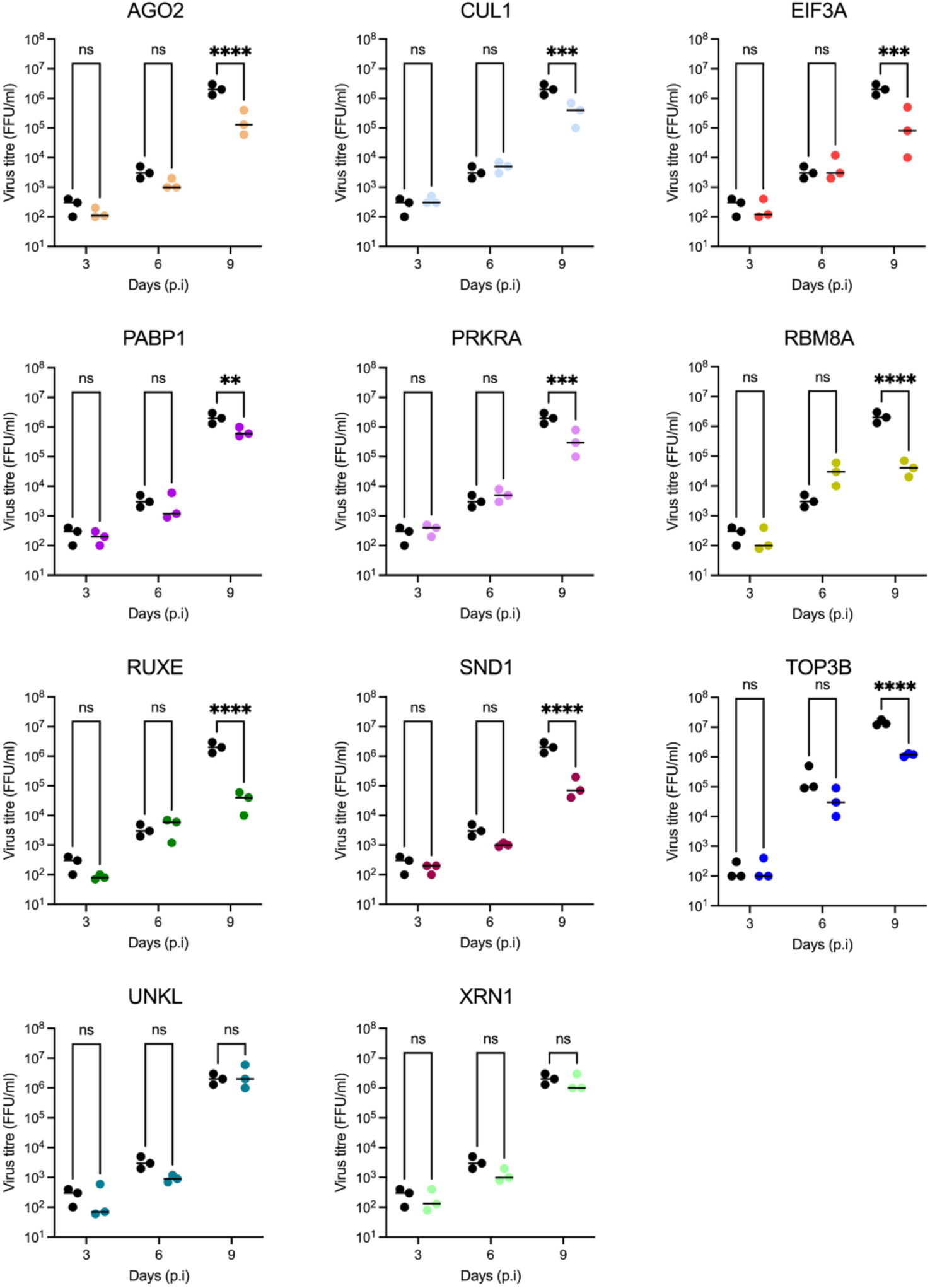
Effect of dsRNA transfections on UUKV titre in supernatant from UUKV-infected ISE6 cell cultures. UUKV titre of supernatant of UUKV-infected, dsRNA-transfected ISE6 cell monolayers, prepared as shown in Fig. 4. Black symbols represent dseGFP data (mock) and coloured symbols represent dsGene data. Statistical significance was measured by ordinary two-way ANOVA with Tukey’s multiple comparisons test. Asterisks indicates significance **** = p < 0.0001, *** = p ≤ 0.001, ** = p ≤ 0.01, * = p ≤ 0.05, ns = not significant.

**Fig 6.**
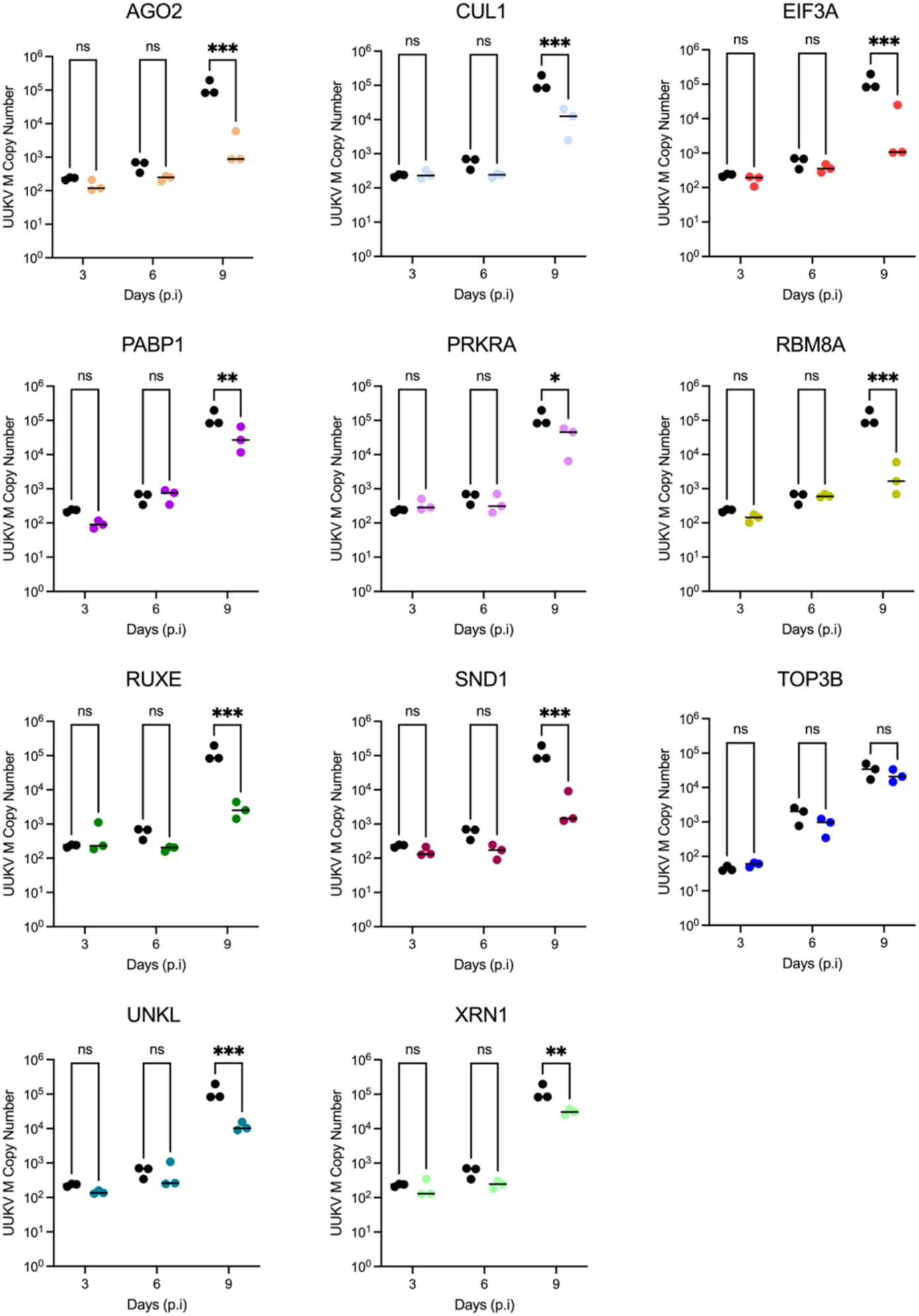
Effect of dsRNA transfections on quantity of UUKV M segment RNA present in supernatant of UUKV-infected ISE6 cell cultures. The copy number of UUKV M RNA within the supernatant of UUKV-infected, dsRNA-transfected ISE6 cell monolayers was determined as prepared as seen in Fig. 4. Black symbols represent dseGFP data (mock) and coloured symbols represent dsGene data. Statistical significance was measured by ordinary two-way ANOVA with Tukey’s multiple comparisons test. Asterisks indicates significance **** = p < 0.0001, *** = p ≤ 0.001, ** = p ≤ 0.01, * = p ≤ 0.05, ns = not significant.

Extracellular viral RNA is a proxy for the total number of released viral particles, including defective non-infectious particles. Both UNKL and XRN1 knockdown caused a decrease in the amount of viral RNA present in the supernatant with no overall effect in viral titre, suggesting that the knockdown of these genes may increase the infective to non-infective particle ratio (Fig 5 & Fig 6). Conversely, TOP3B knockdown caused a decrease in viral titre with no effect on overall extracellular viral RNA levels, which indicates a reduction in viral particle infectivity. Altogether, our data revealed three RBPs (UNKL, XRN1 and TOP3B) with opposite effects on viral particle infectivity.

## DISCUSSION

Research into arboviruses that are transmitted by ticks have largely focussed on the interactions the virus establishes with vertebrate cells and systems. A major contributing factor to the dearth of research into tick-virus interactions has been the lack of characterised cell cultures with corresponding molecular tools, and the difficulties surrounding *in vivo* systems, as previously mentioned. The evolution of high-throughput technology and continuous characterisation of both cell cultures and whole organisms [49] has allowed the genomes of both the *I. scapularis* tick [50] and derived ISE6 cell line [17] to be annotated. From this, researchers have begun to probe into how tickborne arboviruses manipulate the vector cell to facilitate their own replication and transmission.

The advances in tick reagents and resources enabled us to apply comparative RIC to tick cells. The RBPome and its responses have been profiled in several species to date using RIC methodology, including humans [41], mice [51], fish (*Danio rerio*) [52], *Drosophila spp.* [53], parasites (*Leishmania spp.*) [54], yeast (*Saccharomyces cerevisiae*) [55], and plants (*Arabidopsis thaliana*) [56]. In this manuscript we adapted the use of RIC to UUKV-infected ISE6 cell cultures and established a baseline methodology for investigating the effect of virus replication upon the infected tick cell. Despite bunyavirus mRNA molecules lacking a poly-A tail [38,39], the genome of UUKV contained sufficient poly-A like sequences to allow for viral RNA capture.

Therefore, both viral and cellular mRNA were isolated with the oligo(dT) pull down, contributing collectively to the composition of the RBPome. Strikingly, infection of tick cells with UUKV altered the RNA-binding activity of hundreds of RBPs in analogy to what occurs in humans [31,32]. These changes affected different pathways of RNA metabolism and are in line with a tick cell rewiring by two opposite processes: i) the virus activating or inactivating pathways that are required or detrimental (respectively) to infection; and ii) the host cell responding to this cue by triggering the antiviral program illustrated by AGO2, CNOT1 and XRN1. Moreover, a substantial proportion of regulated RBPs interacted with UUKV N in cells, suggesting a direct connection with viral ribonucleoproteins taking over the host RBP machinery and inducing changes to the intracellular RBPome. Several of the RBPs identified as impacting UUKV have pivotal roles in central machineries of RNA metabolism in both vertebrate and invertebrate cells. The phenotypes observed during the loss-of-function experiments could be due to a direct effect on viral RNA, as seen in other studies [57,58], or an indirect effect resulting from the disruption of a central pathway important for cell homeostasis. For example, depletion of the eukaryotic translation initiation factor 3a (eIF3a) and polyadenylate-binding protein 1 (PABP1) were found to increase the amount of UUKV M RNA in the cell at 3 and 6 days p.i. It is unlikely that the activity of PABP1 is on stability or translation of UUKV RNAs because they lack poly(A) tails [38].

Therefore, we hypothesise that the PABP and eIF3a upregulation of viral gene expression is due to an indirect effect by suppressing the translation of cellular mRNAs [59] and increasing the availability of ribosomes for viral mRNAs. Other bunyaviruses such as Rift Valley fever virus (RVFV) utilise viral proteins (NSs) to target PABP1 for degradation in RVFV-infected human cell lines, creating an environment that favours viral translation suggesting a similar mechanism for UUKV in tick cells [60]. While the PABP1 effect can be explained by the lack of poly(A) tail in UUKV RNAs, the effect of the eIF3a knock down is surprising. eIF3a is a core component of the eIF3 complex that bridges the 40S ribosomal subunit and the mRNA through interaction with the eIF4F complex [61]. These interactions are pivotal for initiation of translation of cellular mRNAs, and the fact that UUKV gene expression is favoured by eIF3a implies that UUKV mRNAs follow a non-canonical translation beyond the lack of contribution of PABP. Other viruses encoding internal ribosome entry sites (IRES), such as hepatitis C virus (HCV) [62] and cricket paralysis virus (CrPV) [63] have been shown to have no or partial dependency on eIF3. However, no IRES has been reported at the 5’ leader sequences of UUKV mRNAs, which are derived from the cap-snatching of the cellular mRNAs [64]. How these viral mRNAs translate independently of eIF3a and PABP is unknown and potential explanations such as the existence of a virus-encoded translation factor or the efficient liberation of ribosomes by the virus-induced shutoff should be explored in future. The increase in the level of UUKV RNA following dsRNA knockdown of eIF3a or PABP1, surprisingly, did not lead to an increase in the titre of released virus over the 9-day period. However, this may simply reflect the slow replication kinetics of UUKV within tick cells [2].

Other RBPs tested here are also components of central machineries involved in RNA metabolism, including RBM8A (exon junction complex) [65], RUXE (spliceosome) [66], SND1 (miRNA decay) [67] and XRN1 (mRNA decay) [48]. We observed phenotypes in UUKV infection for all these proteins, but whether they function indirectly through regulation of cellular RNA metabolism or directly through binding to viral RNA, should be elucidated in future work. SND1, for example, could regulate UUKV expression through its influence on the set of RNAis available in the cell. However, recent work has shown that SND1 is critical to recruit NSP9 to the end of SARS-CoV2 RNA to initiate replication [68,69]. Hence, RBPs must be evaluated mechanistically on an individual basis and the indirect versus direct effects should be considered.

Metabolic activity changes could be mediated either by the gene being essential for cell function (exemplified in *CUL1*, *PRKRA*, and *XRN1*) or because increased virus replication observed in some cases (e.g. EIF3A or PRKRA) anticipates viral derived cytopathic effects. This invariably led to a decrease in infectious virus production in knockdown cells by 9 days p.i. One of the most interesting results was that found with the knockdown of *ISCI010954* (ortholog of *TOP3B* in mammalian systems). TOP3B is a topoisomerase that changes the topological state of genetic material through breaking and then reforming genomic nucleic acid strands to unwind supercoils, resolve catenates, and undo knots [70]. This genetic material is usually double-stranded (ds)DNA. However, TOP3B is the only known topoisomerase to interact with both DNA and RNA [71]. TOP3B is dispersed within the cytoplasm and can interact with single stranded nucleic acids through an RGG box [71]. It is also speculated that TOP3B interacts with other RNA-binding proteins to regulate mRNA translation and may also increase the stability of mRNAs [72]. To further support our hypothesis that the ISCI010954 protein has a propensity to bind nucleic acids, we compared the predicted structure of this protein to the mammalian ortholog, TOP3B. To accomplish this, we utilised predicted AlphaFold structures of the human and tick proteins [73]. The *I. scapularis* ISCI010954 (i.e. TOP3B) protein and its *H. sapiens* ortholog show similar spatial arrangement of the topoisomerase-primase (TOPRIM) and DNA topoisomerase type IA domains (Fig. 7A, upper panel), which are essential for TOP3B activity. Moreover, a study employing RBDmap revealed the peptide in the human TOP3B that crosslinks to RNA upon UV irradiation, which represents the RNA-binding surface of the protein. Using *in silico* methods, we determined if this peptide is conserved in *I. scapularis* and found a striking sequence homology (Fig. 7A, lower panel) [74]. This suggests that the human and the tick TOP3B proteins interact with RNA through the same protein-RNA interface. Interestingly, the RNA-bound peptide sequence has higher sequence conservation (around 80%) than the rest of tryptic peptides across the protein sequence (around 60%), suggesting a selective pressure to maintain this sequence unaltered to keep TOP3B function (Fig. 7B).

**Fig 7.**
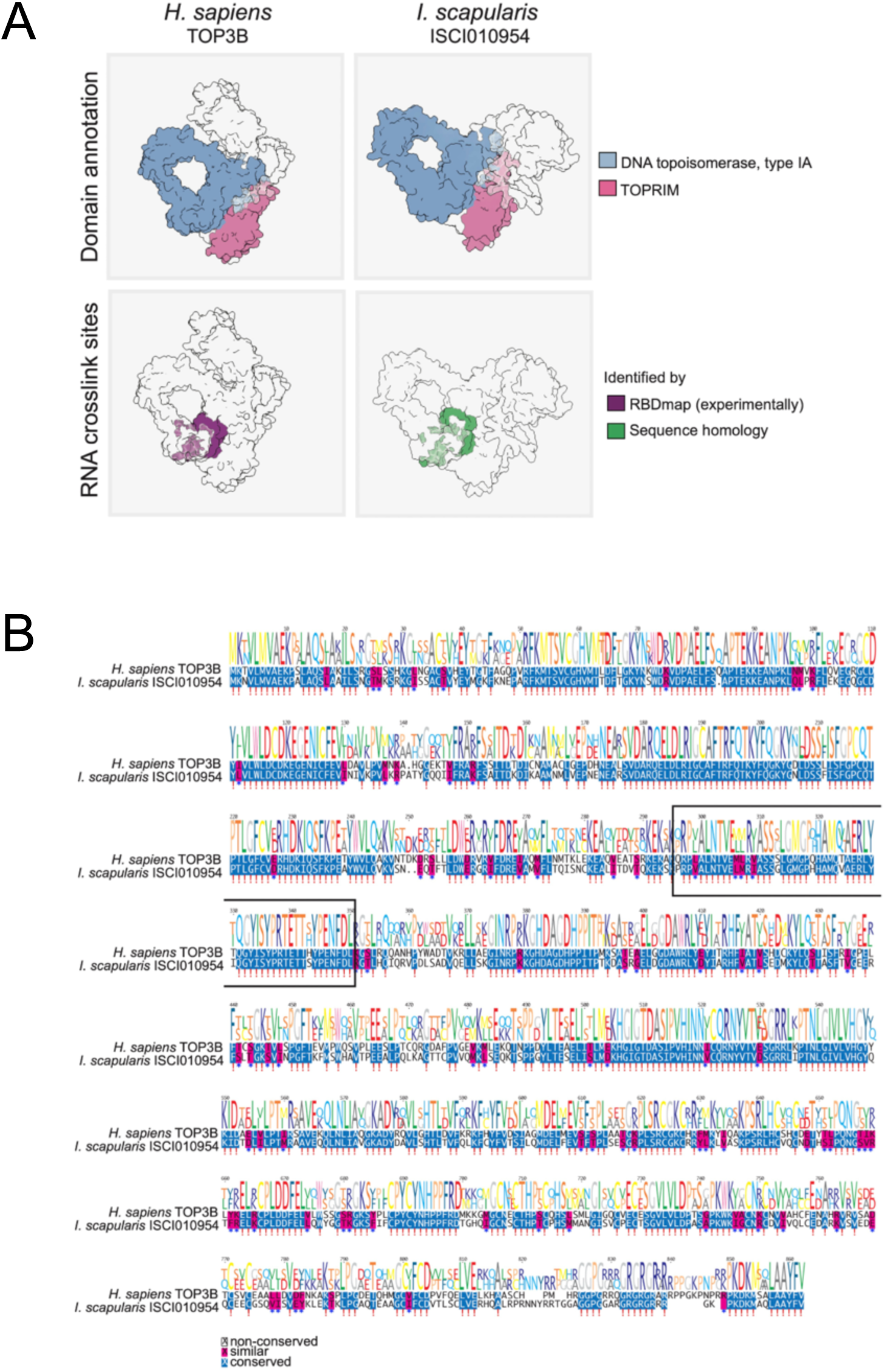
Structure and sequence similarities between *H. sapiens* TOP3B and *I. scapularis* ISCI010954 proteins. (A) AlphaFold structure for *H. sapiens* TOP3B and *I. scapularis* ISCI010954, highlighting the predicted domain annotations (in the upper panel) and the RNA crosslink sites (in the lower panel). (B) Protein sequences alignment of *H. sapiens* TOP3B and *I. scapularis* ISCI010954.

TOP3B was reported to regulate the replication of several flaviviruses, but interestingly it was not reported to affect the replication of two negative sense viruses in mammalian systems [75]. It has been suggested that removal of TOP3B does not affect flavivirus translation or replication, but does impair the production of infectious viral particles, although overall release of virions is unaffected [75]. This is an exciting development as this result was also seen within this study, suggesting that this is potentially a conserved mechanism or function across viruses, and present in both mammals and ticks. The impairment of infectious virus particles following TOP3B depletion maybe be in part due to the function of topoisomerases in ensuring the UUKV RNA is stable and in the correct conformation to be recognised by the N or L proteins and packaged into virus particles. It is unclear if TOP3B itself is packaged into viral particles, however if this is the case it may be used to unwrap the UUKV RNA from the viral nucleocapsid to promote initial viral infection. While most topoisomerases bind DNA and have a nuclear subcellular localisation, under certain conditions, TOP3B can bind mRNAs associated with neuronal cell development in stress granules within the cytoplasmic compartment, and it is also necessary for normal synapse formation in both *D. melanogaster* and mice [76]. Given the predominantly neuronal-like phenotype of the ISE6 cell line [77,78], and the similarities between viral replication factories and stress granules [79] within the cytoplasm of infected cells, it is tempting to speculate that TOP3B is performing a similar function in the infected tick cell and hence impacting virus replication.

Here we focused on the RBPs that were upregulated by UUKV infection, however, downregulated proteins might also be important in virus infection [31,32]. We identified several critical anti-viral RBPs within UUKV infection, which provides the foundation for not only analysis of conserved function across arbovirus species but further investigation into the mechanisms of inhibition which may provide future therapeutic targets.

## ACKNOWLEDGEMENTS

We thank Prof. Ulrike Munderloh, University of Minnesota, for permission to use the ISE6 cell line.

## AUTHORS’ CONTRIBUTIONS

Conceptualization: A.K., A.C., B.B., Data Curation: A.W., W.K., D.L., Y.D., A.C., B.B., Formal Analysis: A.W., W.K., Z.R.deL., R.A., D.L., Y.D., M.N., S.M., A.C., B.B., Funding Acquisition: S.M., A.C., B.B., Investigation: A.W., W.K., K.D., Y.D., M.N., Methodology: A.W., W.K., Z.R.deL., R.A., D.L., E.K., S.M., A.C., B.B., Project administration: A.C., B.B., Resources: A.W., W.K., K.D., L.B.S., M.N., S.M., A.C., B.B., Software: Supervision: A.K., S.M., A.C., B.B., Validation: A.W., W.K., A.C., B.B., Visualization: A.W., W.K., Z.R.deL., R.A., A.C., B.B., Writing – original draft: A.W., W.K., A.C., B.B., Writing – review & editing: A.W., W.K., K.D., L.B.S., D.L., Y.D., E.K., M.N., A.K., S.M., A.C., B.B.

## DATA AVAILABILITY

The data that support the findings of this study are openly available from Enlighten Research Data. Authors will make reagents described in this study available on request (by qualified researchers for their own use). Requests should be directed to the corresponding author.

## MATERIALS AND METHODS

### Experimental model and subject detail

#### Cell culture

Mammalian cell cultures used in this study were BSR and BSR-T7/5-CL21 (referred to as BSR-T7) cells, provided by Karl-Klaus Conzelmann of Ludwig-Maximilians-Universität München. The BSR cell line is a clone of BHK-21 [80]. BSR-T7 cell cultures are a modified BSR cell line stably transfected with the bacteriophage T7 RNA polymerase [81]. Both cell lines were maintained in DMEM (Thermo Fisher Scientific) supplemented with 10% (v/v) foetal bovine serum (FBS) at 37°C in an atmosphere of 5% CO_2_ in air. BSR-T7 cell medium was supplemented with 1mg/ml of the selection agent Geneticin (G418) prior to transfection. The tick cell line used in this study was ISE6, derived from embryonic *I. scapularis* [35], sourced from the Tick Cell Biobank at the University of Liverpool. The cells were grown in sealed, flat-sided tubes (Nunc) for maintenance and transferred to sealed T25 or T80 non-vented flasks to bulk them up for experiments. ISE6 cells were maintained at 32°C in enriched L-15B300 medium [82], supplemented with 10% tryptose phosphate broth, 5% FBS, 2 mM l-glutamine and 0.1% bovine lipoprotein concentrate (MP Biomedicals, Thermo-Fisher).

#### Virus Production

UUKV used in this project was a recombinant UUKV generated using reverse genetic technologies and based upon the prototype tick isolate S-23 [3]. Working stocks of UUKV were generated from recombinant virus rescued in BSR-T7 cells, followed by amplification and generation of working stocks in BSR cell cultures. UUKV stocks were titrated using an immunofocus assay as described below.

#### Immunofocus Assay

Virus titres were enumerated by an immunofocus assay in BSR cells. Briefly, confluent monolayers of BSR cells were infected with serial dilutions of virus made in phosphate buffer saline (PBS) containing 2% FBS and incubated for 1h at 37 °C, followed by the addition of a GMEM overlay supplemented with 2% FCS and 0.6% Avicel (FMC Biopolymer). The cells were incubated for 5 days before fixation and subsequent use in focus-forming assays as described previously [3,14].

### RNA interactome capture (RIC)

Comparative RNA interactome capture (RIC) was performed following a previously described protocol [34], with the following modifications. ISE6 cells were seeded in six sets of 3 x 10cm plates with 1.5 x 10^7^ cells/plate. Cells were either mock-infected or inoculated with UUKV at a multiplicity of infection (MOI) of 5 FFU/cell. Samples were harvested from three replicate plates at 9 days p.i., when cells were washed 3x with PBS, then irradiated with 150 mJ/cm^2^ of UV light at 254nm on ice and lysed with 3ml of lysis buffer (20mM Tris-HCl [pH 7.5], 500mM LiCl, 0.5% [w/v] LiDS, 1mM EDTA, 0.1% [v/v] IGPAL, and 5mM DTT) Lysates were then mechanically homogenised by passing the sample through a 32G diameter needle using a 5ml syringe on ice. A sample of homogenized lysate was removed at this stage for silver staining and qPCR analysis. Oligo(dT)_25_ capture beads were equilibrated in lysis buffer, and 300µl of the vortexed bead slurry mix was added to each of the 6 samples and incubated for 1 h at 4°C with gentle rocking. Beads were collected in 1.5ml microcentrifuge tubes via a magnet, and the lysates removed and discarded. Beads were then washed as described below with lysis buffer, wash buffer 1 (20mM Tris-HCl [pH 7.5], 500mM LiCl, 1mM EDTA, 0.01% [v/v] IGEPAL, and 5mM DTT), wash buffer 2 (20mM Tris-HCl [pH 7.5], 500mM LiCl, 1mM EDTA, 0.01% [v/v] IGPAL, and 5mM DTT) on ice, followed by wash buffer 3 (Oligo(dT) buffer 3: 20mM Tris0HCl [pH 7.5], 200mM LiCl, 1mM EDTA, and 5mM DTT) at room temperature. To wash, 1ml of the appropriate buffer was added to the beads. The beads were inverted 10 times per minute, for five minutes, whilst being stored on ice or at room temperature as specified. The beads were washed 3 times with each of the buffers utilised in the protocol. Beads were resuspended in 125μl of elution buffer (20mM Tris-HCl [pH 7.5] and 1mM EDTA) and incubated for 3 min at 55°C with agitation. Finally, prepared eluates were stored at -80°C.

### Coimmunoprecipitation

Coimmunoprecipitation (co-IP) was carried out using ISE6 cell monolayers infected with UUKV at a MOI of 5 FFU/cell, as described above. Mouse anti-UUKV N antibodies, generated from hybridoma 8B11A3 (kindly provided by Dr Anna Överby Wernstedt, Umeå University [83]) were cross-linked to Dynabeads™ Protein G (Thermo Fisher) using BS3 (bis(sulfosuccinimidyl)suberate) following the manufacturer’s instructions. Antibody cross-linked beads were prepared for use in immunoprecipitation by washing in 500µl of lysis buffer (20mM Tris, 150mM NaCl, 5mM MgCl2, 0.5% [v/v] NP40, 25 unit/ml Benzonase (Merck Millipore), fresh cOmplete protease inhibitor [1 tablet per 10ml of buffer], phosphatase inhibitor [Roche, 1 tablet per 10ml of buffer] per 100µl bead mix with gentle rotation at 4°C for five min and finally stored in lysis buffer at 4°C.At 9 days p.i., cells were lysed in 1ml of lysis buffer per plate. Cell lysates from either mock or UUKV-infected triplicate plates were pooled and kept on ice.

Supernatants were clarified by centrifugation for 20 min at 5000 rpm and 4°C and transferred to fresh 15 ml centrifuge tubes where 100µl of the lysates were removed for analysis by silver staining. Prepared antibody-cross-linked beads were incubated with the cleared lysates and incubated with gentle rotation at 4°C overnight. Beads were then collected using a magnet, and the supernatant discarded. Beads were washed with 500µl of wash buffer (50mM Tris, 200mM NaCl, 1mM EDTA, 1% [v/v] NP40, fresh cOmplete protease inhibitor [1 tablet per 10ml / buffer]) with gentle rotation at 4°C for five min. Washing was repeated three times then the proteins were eluted by incubation of the beads with 125µl of elution buffer (100mM TEAB, 5% SDS [w/v]) at 95°C for 10 min. Protein eluates were stored at -80°C prior to subsequent analysis.

### Synthesis of double-stranded RNA (dsRNA)

RNA was extracted from ISE6 cells using TRIzol (Invitrogen), following the chloroform extraction protocol. Reverse transcription (RT) was carried out using the SuperScript™ Reverse Transcriptase kit (Invitrogen) and random hexamers (50µM) to produce cDNAs corresponding to cellular or viral targets (listed in Table S1). Unique portions of the gene candidates were amplified from the prepared cDNAs with primers incorporating a minimal T7 RNA polymerase promoter sequence (All primer sequences can be found in Table S2). PCR products were analysed by agarose gel electrophoresis and purified by gel extraction. PCR products were sequenced for target verification before the production of dsRNA. The MEGAscript® RNAi Kit (Thermo Fisher Scientific) was used to produce the dsRNA from the PCR fragments, according to the manufacturer’s instructions. To determine the quality of the dsRNA, the concentration was determined using a Nanodrop One. Once prepared dsRNAs were stored at -80°C.

### dsRNA knockdown in tick cells

ISE6 cells were seeded into 24-well plates and monolayers were transfected using the Magnetofectamine O2 transfection kit (1ug of dsRNA/1x10^6^ cells), at the indicated ratio of dsRNA:transfection reagent, 2µl of magnetofectamine beads, and 250µl OptiMEM (Gibco), following the manufacturer’s protocol (Oz Biosciences). After transfection, the transfection medium was removed, and fresh medium applied. Monolayers were then incubated for 20 h before infection with UUKV at a MOI of 5 FFU/cell. Cell monolayers were harvested at the time points indicated and analysed using RT qPCR, cell metabolic activity assays, whole cell immunofluorescence analysis, and focus-forming assay to assess the production of infectious virus particles.

### Whole cell immunofluorescence analysis

For immunofluorescence analysis of UUKV-infected cultures, culture supernatants were removed, and monolayers washed with PBS. Cells were fixed in 8% formaldehyde solution for 1 h and permeabilised in permeabilization buffer (0.5% [v/v] Triton X-100 [Roth] in PBS) for 30 min. For both primary and secondary antibody buffers, the corresponding antibodies (Mouse anti-UUKV generated from hybridoma 8B11A3 and Anti-mouse IgG (H&L) Secondary Invitrogen #T6199, respectively) were diluted in blocking buffer (4% (w/v) skimmed milk powder (Marvel) in PBST - 0.1% (v/v) Tween [Sigma] in PBS). The cell monolayers were then washed three times with PBS before being probed with primary antibody buffer. Primary antibody incubation was carried out overnight at 4°C. Cells were then washed three times with PBS before being probed for 1 h at room temperature with secondary antibody. Cells were washed once with PBS before being incubated with DAPI diluted in PBS (3µl:10mL buffer) for 10 min at room temperature. Finally, cell monolayers were washed three times with PBS. Cells were kept at 4°C in PBS until imaging. Monolayers were imaged using an Odyssey® CLx Imaging System (Li-Cor) and analysed using the associated software to determine the total UUKV N fluorescence compared to a mock-infected well as a control.

### Cell metabolic activity assay

CellTiter-Glo® 2.0 Cell Viability Assay (Promega) was used to determine the metabolic activity of transfected cell monolayers. At the appropriate timepoint, cell monolayers were washed before being resuspended in 150µl of fresh PBS. 50µl of the cell suspension or PBS control was pipetted into opaque-walled 94 well plates in triplicates. The cell suspension was then mixed with 50µl of viability reagent and incubated in the dark for 10 min at room temperature. Well luminescence was measured using the GloMax® Navigator Microplate Luminometer (Promega).

### Reverse-transcription and quantitative PCR

RNA was isolated from cell monolayers or infected cell supernatant using TRIzol, following the chloroform extraction protocol (Invitrogen). Reverse transcription (RT) to generate cDNA was carried out using the SuperScript™ Reverse Transcriptase kit and random hexamers (50µM; Thermo Fisher Scientific), following the manufacturer’s instructions. Positive controls were generated from randomly primed cDNA derived from either total cell RNA or UUKV-infected cell culture supernatant. For negative controls, cDNA was replaced with nuclease-free H2O. qPCR primers for UUKV were designed to target the genomic M segment RNA. qPCR analysis was undertaken using SYBR-green (Applied Biosystems) on a QuantStudio 5 Real-Time PCR System (Applied Biosystems) and associated software. Details of the primers used for qPCR in this study can be found in Table S2. Transcript expression levels relative to the ISE6 18S ribosomal subunit as a reference were calculated according to the 2^-ΔΔ*CT*^ methodology [84]. Where no housekeeping gene was present to allow for the ΔΔCt calculation, such as when isolating UUKV M RNA from supernatant, Ct values were normalized, or a standard curve was produced to allow gene copy number to be determined. The maximum detection limit for the thermocycler was defined as a Ct of 40, and negative controls produced a Ct of 30 or above. Therefore, any samples which produced a normalised Ct ≤ 10 were therefore classified as a negative result.

### Conventional Protein Analysis

If cell monolayers were not lysed through prior protocols, lysis of mock or UUKV-infected cell monolayers was carried out using Laemmli buffer (100mM Tris-HCl, 4% (v/v) SDS, 20% (v/v) glycerol, 200mM DTT, 0.2% bromophenol blue (v/v), 3µl/ml endonuclease). Samples were resolved on SDS-PAGE and visualised via Pierce™ Silver Stain Kit (Thermo Fisher Scientific #24612) and/or by western blotting using mouse anti-UUKV N and anti-mouse IgG (H&L) secondary, the Li-Cor Odyssey system for visualization and the Image Studio Lite software (Li-Cor) for quantification. Data shown in the manuscript are representative gels from at least three independent replicates.

### Mass spectrometry and relative protein quantification

RIC and viral nucleocapsid protein (NCAP) pulldown eluates were processed through single-pot solid-phase-enhanced (SP3) as previously described [41] and SDS-PAGE followed by in-gel digestion sample preparation, respectively. After SP3 processing, prepared samples of ISE6 RIC were analysed at the Rosalind Franklin Institute. Analysis of peptides was carried out using an Ultimate 3000 nano-LC 1000 system coupled to an Orbitrap Fusion Lumos (Thermo Fisher Scientific). Samples were diluted in ultra-pure water with 5% formic acid and 5% DMSO prior to injection. Peptides were initially trapped on a C18 PepMap100 pre-column (300 µm inner diameter x 5 mm, 100A) and then separated on an in-house built C18 column (Reprosil-Gold, Dr. Maisch, 1.9 μm particle size) column (ID: 50 μm, length: 50 cm) at a flow rate of 100 nL/min. Peptides were separated over 60 min using mobile phase A (water and 5% DMSO, 0.1% formic acid) and a 12-30% gradient of mobile phase B (acetonitrile and 5% DMSO, 0.1% formic acid). Separated peptides were directly electro-sprayed into an Orbitrap Fusion Lumos mass spectrometer (Thermo Fisher Scientific) and analysed in a data-dependent mode. MS1 spectra were acquired in the orbitrap (350-1400 m/z, resolution 60000, AGC target 1.2 x 10^6, maximum injection time 50 ms). The top 40 most abundant peaks in the survey scan were fragmented using HCD and analysed in the ion trap (scan speed Turbo, AGC target 1 x10^4, maximum injection time 32 ms, normalised collision energy 30%). Nucleocapsid protein pulldown eluates were sent to the Dundee FingerPrints Proteomics facility for mass-spectrometry analysis (in data-dependent mode and label-free quantification). Samples were resolved via 1D SDS-PAGE on a 10% gel with MOPS buffer and stained with Quick Coomassie Stain (Generon). Each gel lane was run for 15 min with the whole area being excised, then subjected to in-gel digestion using 1mg/ml Trypsin (Thermo) at a final concentration of 12.5µg/mL. Digested peptides were run on a Q-Exactive HF (Thermo Scientific) instrument coupled to a Dionex Ultimate 3000 HPLC system (Thermo Scientific). A 2-35%B gradient comprising of eluent A (0.1% formic acid) and eluent B (80% acetonitrile/0.1% formic acid) was used to run a 120-minute gradient per sample. The top 20 most intense peaks from a mass range of 335-1800 m/z in each MS1 scan with a resolution of 60,000 were then taken for MS2 analysis at a resolution of 15,000. Spectra were fragmented using Higher-energy C-trap dissociation (HCD). The raw data files were provided upon completion for both sets of eluates. Protein identification and quantification were obtained using the Andromeda search engine implemented in MaxQuant (v2.4.11.0) and searched against the ISE6 reference proteome [19] and UUKV proteome [28] with ‘match between run’ activated and under default parameters. MaxQuant outputs (proteinGroups) were used for downstream relative quantification. Potential protein contaminants flagged by MaxQuant were filtered out together with proteins with missing values across all samples, using R-package “DEP (1.4.1)”,. For relative quantification between UV-crosslinked versus non-crosslinked infected samples, protein raw intensities were log_2_-transformed. Missing value imputation was performed only for proteins undetected in all replicates in one experimental condition, while present in the other condition (in at least 2 replicates). Imputation was implemented using minimum determination method (Mindet) ^64^. Next, the processed protein intensities were subjected to statistical analysis using the empirical Bayesian method moderated t-test with p-values adjusted for multiple-testing (Benjamini-Hochberg method) provided by R-package “limma (3.38.3)” for. For all other comparisons, protein intensities were processed as described above with inclusion of a normalization step for raw protein intensities using R-package Variance Stabilizing Normalization “VSN (3.50.0)”.

### Protein ortholog identification

To identify human (Uniprot_id: UP000005640, downloaded Nov2016) and fly (Uniprot_id: UP000000803, downloaded Aug2024) orthologs of ISE6 proteins InParanoid-DIAMOND was used under default settings [85,86].

### Reactome enrichment analysis

Enrichment analysis of cellular pathways in differentially regulated RNA binding proteins was performed by R package ReactomePA. The analysis only included ISE6 proteins with human orthologs and all human genes as background [87].

### Structure prediction, protein domains and RNA crosslink Sites annotation

Using ColabFold [88] in batch mode we predicted protein structures. We exported the “relaxed_rank_001_alphafold2” pdb file to Chimerax v1.7 for visualization [89]. Protein domains identified by InterProScan [90] and RNA crosslink sites identified by RBDmap [74] or by sequence homology by ClustalOmega [91].

### Quantification and Statistical Analysis

For each condition, biological triplicates were produced, where each biological sample was tested (for example by RT qPCR) in triplicate, and an average of the technical replicates was then used for plotting and statistical analysis. Testing for statistical significance was carried out by either unpaired t-test, one-way ANOVA, or two-way ANOVA with Tukey’s multiple comparison. The cut-off for significance was set at p<0.05. Where analysis was employed, unless stated otherwise both “ns” and a lack of notation indicate no significance.

