## Supplementary Information for "Uukuniemi virus infection causes a pervasive remodelling of the RNA-binding proteome in tick cells"

**The PDF file includes:**

**Fig S1.** Diagnostics of mass spectrometry results from RIC biological triplicates and immunoprecipitation biological triplicates, and further analysis of immunoprecipitation mass spectrometry results.
**Fig S2.** Effect of dsRNA transfections on selected gene expression within ISE6 cell culture.
**Fig S3.** Effect of dsRNA transfections on the UUKV N protein expression within UUKV infected ISE6 cell culture.
**Table S1.** Table of oligonucleotide primers used in the production of double stranded RNA, including primer name, sequence and description of use.
**Table S2.** Table of oligonucleotide primers used in the production of double stranded RNA, including primer name, sequence and description of use.

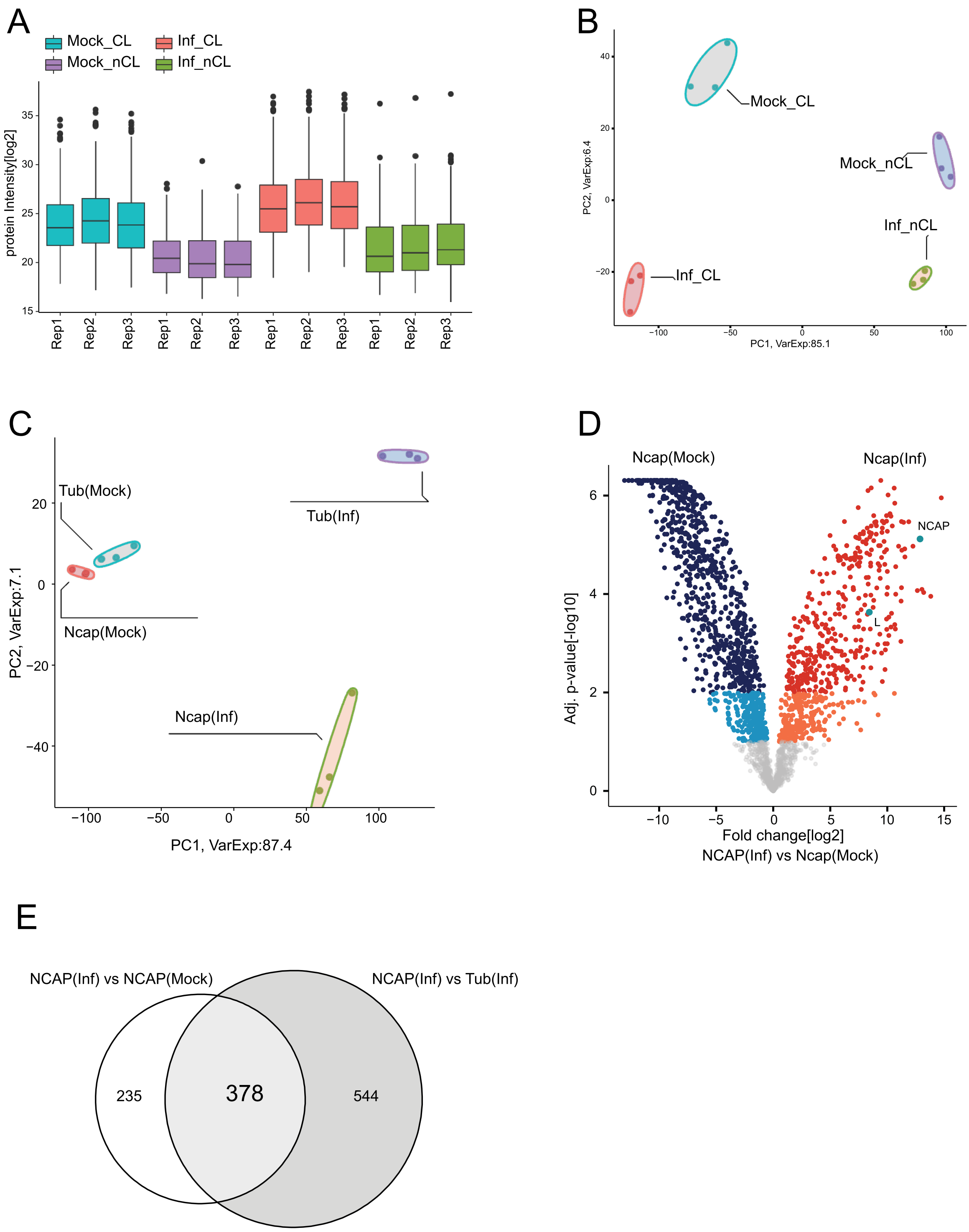


**Fig S1. Diagnostics of mass spectrometry results from RIC biological triplicates and immunoprecipitation biological triplicates, and further analysis of immunoprecipitation mass spectrometry results.**
Diagnostics of mass spectrometry data derived from rUUKV-infected cRIC eluates as described in Fig 1 and mock or rUUKV-infected immunoprecipitation eluates as described in Fig 2.
**(A)** The protein intensities (intensity[log2], y axis) for all proteins found within each triplicate sample (x axis) of each condition; mock or rUUKV infected, crosslinked or non-crosslinked, were plotted from rUUKV-infected cRIC eluates.
**(B)** Principal component analysis (PCA) was performed from rUUKV-infected cRIC eluates. The plot models 91.5% of the total data variance. Variance proportions are shown along both axes, and groupings of the triplicates for each sample are highlighted on the plot.

**(C)** Principal component analysis (PCA) was performed for mass spectrometry results of mock or rUUKV-infected immunoprecipitation eluates. The plot models 94.1% of the total data variance. Variance proportions are shown along both axes, and groupings of the triplicates for each sample are highlighted on the plot.

**(D)** The fold change in the proteins specific to UUKV N immunoprecipitation in mock- and r-UUKV-infected cells (x axis) were plotted against the significance of this fold change (Adj. p value[-log10], y axis). Proteins with a positive fold change, 1-10% FDR (orange), ≤1% (red); proteins with a negative fold change, 1-10% FDR (light blue), ≤1% (dark blue) and proteins with greater than 10% FDR are shown in grey.

**(E)** The infected cell tubulin vs UUKV N protein pulldown (Fig. 2F) and mock infected vs UUKV infected UUKV N pulldown (Fig. S1D) was cross-referenced to identify proteins within both groups, of which 378 proteins were found to be present in both groups.

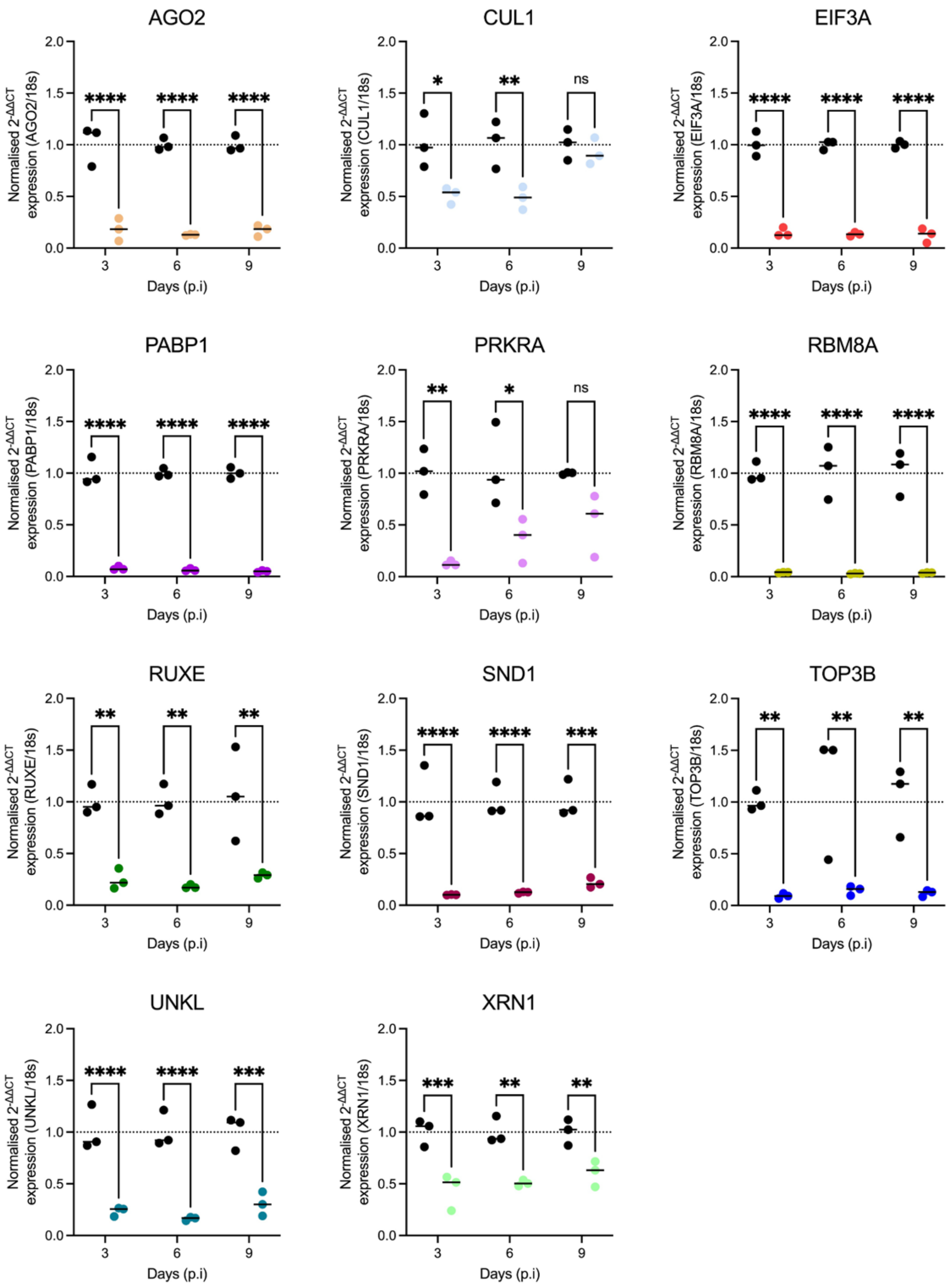


**Fig S2. Effect of dsRNA transfections on selected gene expression within ISE6 cell culture.**

Triplicate ISE6 cell monolayers were transfected with 2µg of dsRNA homologous to the indicated gene. Black symbols represent dseGFP data (mock) and coloured symbols represent dsGene data. At one day post-transfection, cell monolayers were infected with rUUKV at 5 FFU/cell. At timepoints indicated, cell monolayers were lysed via TRIzol and cellular RNA extracted. RT qPCR was performed on the cellular RNA using primers against the indicated gene, and I. scapularis ribosomal 18S as the ‘housekeeping’ gene. The normalised expression of UUKV M RNA was calculated using the 2-ΔΔCt method. Statistical significance was measured by ordinary two-way

ANOVA with Tukey's multiple comparisons test. Asterisks indicates significance **** = p < 0.0001, *** = p ≤ 0.001, ** = p ≤ 0.01, * = p ≤ 0.05, ns = not significant.

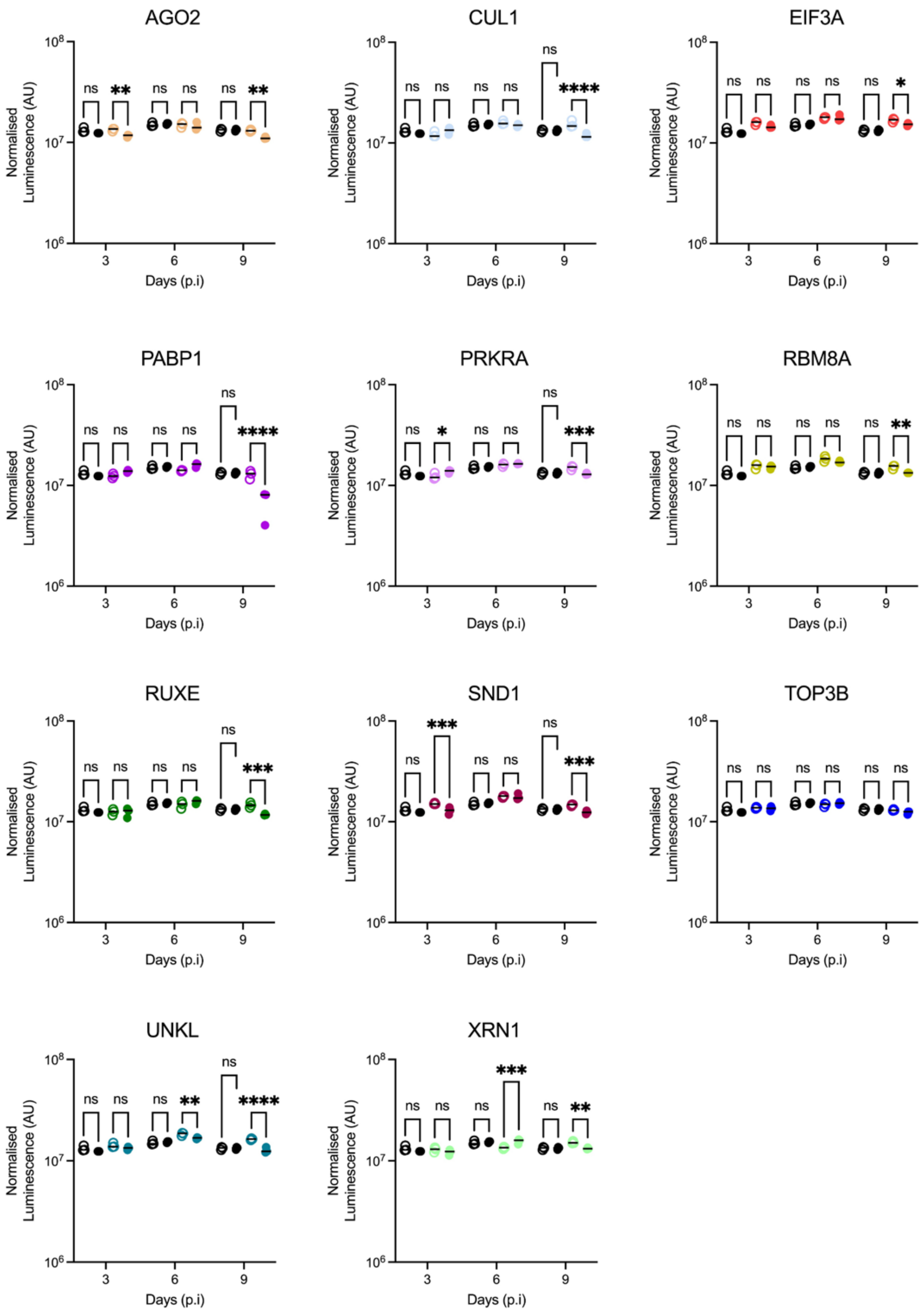


**Fig S3. Effect of dsRNA transfections on the UUKV N protein expression within UUKV infected ISE6 cell culture.**

Triplicate ISE6 cell monolayers were transfected with 2µg of dsRNA homologous to the indicated gene. Black symbols represent dseGFP data (mock) and coloured symbols represent dsGene data. At one day post-transfection, cell monolayers were infected with rUUKV at 5 FFU/cell. At timepoints indicated, cell monolayers were fixed using formaldehyde. Cell monolayers were then probed using mouse anti-UUKV primary antibody and antimouse fluorescent antibody. The fluorescent intensity of each sample was normalised against non-infected cells. Statistical significance was measured by ordinary two-way ANOVA with Tukey's multiple comparisons test. Asterisks indicates significance **** = $p < 0.0001$, *** = $p \le 0.001$, ** = $p \le 0.01$, * = $p \le 0.05$, ns = not significant.

**Table S1: Table of oligonucleotide primers used in the production of double stranded RNA, including primer name, sequence and description of use.**

| Primer name | Primer sequence (5' to 3') |
|---|---|
| eGFP dsRNA Forward | GTAATACGACTCACTATAGGGATGGTGAGCAAGGGCGAGGAGCTGTTC |
| eGFP dsRNA Reverse | GTAATACGACTCACTATAGGGCTGGGTGCTCAGGTAGTGGTTGTCGGGC |
| UUKV N dsRNA Forward | GTAATACGACTCACTATAGGGATGAGACCCTCCCTGAGGAC |
| UUKV N dsRNA Reverse | GTAATACGACTCACTATAGGGATCTGAGGACAGTTGCAGCC |
| AGO2 dsRNA Forward | GTAATACGACTCACTATAGGGACGTGAACAAGACGTCTCCC |
| AGO2 dsRNA Reverse | GTAATACGACTCACTATAGGGGGAGCTCCTTCACCATCGAG |
| PABP1 dsRNA Forward | GTAATACGACTCACTATAGGGGGCTGTTCCCCCTCATCCAC |
| PABP1 dsRNA Reverse | GTAATACGACTCACTATAGGGTCACTCCTTCTTGAGCGAG |
| XRN1 dsRNA Forward | GTAATACGACTCACTATAGGGCAACTGCCGGAAAGGTGTTG |
| XRN1 dsRNA Reverse | GTAATACGACTCACTATAGGGCTGCTTGTTGGGTGGCTTTC |
| TOP3B dsRNA Forward | GTAATACGACTCACTATAGGGCCGTCTACGAGTACATGGGC |
| TOP3B dsRNA Reverse | GTAATACGACTCACTATAGGGGGTAGTCACAGCCTTGACCC |
| SND1 dsRNA Forward | GTAATACGACTCACTATAGGGTTGACTACGGCAATCGGGAC |
| SND1 dsRNA Reverse | GTAATACGACTCACTATAGGGGACCAGCAGGGTCACAAAGT |
| RBM8A dsRNA Forward | GTAATACGACTCACTATAGGGGGAAGGCTGGATCCTGTACG |
| RBM8A dsRNA Reverse | GTAATACGACTCACTATAGGGTGCGGCGATGACTTCTTTTC |
| EIF3A dsRNA Forward | GTAATACGACTCACTATAGGGTGCCACACCGTTCTATCTCG |
| EIF3A dsRNA Reverse | GTAATACGACTCACTATAGGGATTCCTTGATCTCGTCCGGC |
| UNKL dsRNA Forward | GTAATACGACTCACTATAGGGCCTGTACGAGTACCAGGGGG |
| UNKL dsRNA Reverse | GTAATACGACTCACTATAGGGTCCAGGTCCTGTGGCCTAA |
| RUXE dsRNA Forward | GTAATACGACTCACTATAGGGGGACCAGGCCAAAAAGTTCAG |
| RUXE dsRNA Reverse | GTAATACGACTCACTATAGGGCCAAACCTGAATCCGAGCCC |
| CUL1 dsRNA Forward | GTAATACGACTCACTATAGGGATGTGCTGCGGTTCTACACA |
| CUL1 dsRNA Reverse | GTAATACGACTCACTATAGGGATCTCGTAGATGCCCTTGCG |
| PRKRA dsRNA Forward | GTAATACGACTCACTATAGGGCTACATGGGGCTGAAGGAGC |
| PRKRA dsRNA Reverse | GTAATACGACTCACTATAGGGGTGCTGAGCTCGTCTATGGG |

**Table S2. Table of oligonucleotide primers used in the production of double stranded RNA, including primer name, sequence and description of use.**

| Primer name | Primer sequence (5' to 3') |
|---|---|
| UUKM Standard Forward | ACTTGGCATCTGCCACCATGTTAATC |
| UUKM Standard Reverse | GCCGACCCACACAAAGACA |
| UUKM qPCR Forward | TGCTACTTTCGGTGCCCTAA |
| UUKM qPCR Reverse | CAGGAGGCTTTGAACCAACC |
| 18s Standard Forward | CGTAGTTCCGACCATAAACGA |
| 18s Standard Reverse | CATCTAAGGGCATCACAGACC |
| 18s qPCR Forward | GACTCAACACGGGAAACCTC |
| 18s qPCR Reverse | TAACCAGACAAATCGCTCCAC |
| AGO2 qPCR Forward | CGAGAGCGGGAGATCAACAA |
| AGO2 qPCR Reverse | GAATGCGACCTCGTACCTCC |
| PABP1 qPCR Forward | ACATGATCACTCGCCGATCC |
| PABP1 qPCR Reverse | TCGGCTTGTTCTTGATGGCA |
| XRN1 qPCR Forward | GCTCCGAATCTCTGGACGAG |
| XRN1 qPCR Reverse | CGCCCGAAAAAGTGACTTGG |
| TOP3B qPCR Forward | GCGTGGAGGCTGTACGATTA |
| TOP3B qPCR Reverse | CCCGGATTGATGACGCTCTT |
| SND1 qPCR Forward | GACAACGGTCACTGGAGGTT |
| SND1 qPCR Reverse | GCGGAGTAGTCCTTCCACAG |
| RBM8A qPCR Forward | CTGAAGGAACGAGCTCGGAA |
| RBM8A qPCR Reverse | TTCATCCGTGTCCATGGCTT |
| EIF3A qPCR Forward | ACCCCCTGGAAGAGAGGTG |
| EIF3A qPCR Reverse | ATGTCCCTCGAACGGTCCAT |
| UNKL qPCR Forward | TACGACGAGACGACGGGTAT |
| UNKL qPCR Reverse | GCCCGTCTTGTAGTAACGCA |
| RUXE qPCR Forward | CTACAGAACAGGGCTCGGATT |
| RUXE qPCR Reverse | GCTTCCGCTGCTTAGTCTTTG |
| CUL1 qPCR Forward | GAGGCCAGCATGATCTCCAA |
| CUL1 qPCR Reverse | AAGGGCCTCTTCGGAGTTTG |
| PRKRA qPCR Forward | TTTGGCATCTCTTGCCTCGT |
| PRKRA qPCR Reverse | ATCATCTTGTAGGCAGCCTGG |
